# DockFormer: Affinity Prediction and Flexible Docking with Pair Transformer

**DOI:** 10.1101/2024.11.25.625135

**Authors:** Ben Shor, Dina Schneidman-Duhovny

## Abstract

Protein-small molecule interactions, or receptor-ligand interactions, are essential for understanding biological processes and advancing drug design. Despite advancements, existing prediction models of these interactions still lack capabilities and accuracy needed to replace traditional screening. In this paper, we introduce DockFormer, a method that leverages multi-modal learning to predict both the binding affinity and structure of these interactions. DockFormer employs fully flexible docking, where no part of the receptor remains rigid, by adapting the AlphaFold2 architecture. Instead of relying on protein sequences and Multiple Sequence Alignments, DockFormer uses predicted receptor structures as input. This modification enables the model to concentrate on ligand docking prediction rather than protein folding, while preserving full receptor flexibility. The streamlined design also reduces the model size to just 8 layers, compared to AlphaFold2’s 48 layers, greatly accelerating the inference process and making it more efficient for large-scale screening. When evaluated on affinity benchmarks such as CASF-2016, PLINDER, and the recently released CASP16 ligand screening benchmark, DockFormer performs comparably to or better than state-of-the-art methods, which typically rely on templates or bound structures as input. On structural benchmarks such as Posebusters and PLINDER, DockFormer demonstrated success rate of 20% and 15%, respectively.

## 1 Introduction

Interactions between proteins and small molecules, or receptor-ligand interactions, are critical for biological functions and drug design[1–3]. A major challenge is the screening process, in which a huge database is searched for ligands that bind to a specific target receptor with high affinity[4]. Currently, this screening process heavily relies on wet-lab experiments and physics-based computations [5], both of which are expensive and time-consuming. The importance of these interactions stresses a need for accurate *in silico* screening prediction methods.

The interaction prediction task is defined as the prediction of structure, prediction of affinity, or both. The prediction of bound interaction structure given the receptor holo- or apostructure is generally known as the docking of a ligand in the receptor. Docking methods can be physics-based, such as GLIDE [6] and AutoDock [7] or deep-learning-based[8, 9], such as DiffDock [10]. These methods use the receptor backbone structure as rigid and change the position and conformation of the ligand and possibly the side-chains. However, the performance of these methods deteriorates if only apo structure is available. Prediction of both protein structure and the interaction is commonly referred to as co-folding. Co-folding methods, while potentially more accurate, are resource-intensive and complex [11, 12]. For many years, a major obstacle to co-folding methods was the lack of accurate methods to predict individual protein structures.

The release of AlphaFold2 [13] marked a significant advancement in protein structure prediction, utilizing a transformer-based architecture called EvoFormer to process single and pairwise sequence representations. In its successor, AlphaFold3, a more lightweight variation named Pairformer was used [11], where Multiple Sequence Alignments (MSA) are preprocessed separately. Although the success of AlphaFold2 was partly due to MSA, studies have shown that even without MSA, the model can predict protein interaction interfaces [14, 15], stressing the competence of this architecture. Despite these advances, structural models generated are not ideal for docking ligands, as evidenced by the reduced accuracy of methods like GLIDE (44% to 15%) and DiffDock(38% to 21%) when using AlphaFold2 predictions [5, 16]. This is because small-molecule binding can alter the protein conformation in mechanisms known as induced fit or population shift [17, 18]. Change can be small, for example, only in the side-chains, or significant when binding to the cryptic allosteric sites [19].

In addition to structure prediction methods, there are methods designed to predict affinity given the bound structure[20–25]. There are also approaches for predicting affinity without the bound structure[26, 27]. These methods can work either with the protein structure or just its sequences. While they perform well when trained on a specific protein, they struggle to generalize to new proteins outside their training set. This limitation often arises because, without the bound structure, the models cannot learn the specific interactions influencing affinity and instead tend to memorize small molecule features and their impact on affinity in specific proteins [28].

Multi-modal learning refers to a machine learning approach where a model processes and integrates multiple types of data (modalities) to improve learning and prediction and previously had some validation for receptor-ligand prediction tasks[29, 30]. Here we apply multi-modal learning to train a multi-task model, DockFormer, that performs two prediction tasks of different data types - structure and affinity. In computer vision, it has been shown that if tasks are similar enough we could get improved accuracy in all tasks by using a single model, as opposed to different specialized models[31]. In the field of protein folding, multi-modal methods such as ESM3[32], have also been proven to be successful in obtaining state-of-the-art performance on multiple tasks such as structure prediction, structure design, and masked position prediction.

Here, we introduce DockFormer, a multi-modal model designed to predict both receptor-ligand affinity and structure. DockFormer leverages the Pairformer architecture to identify biological and physical interfaces without relying on MSAs. Instead, it predicts receptor-ligand contacts based solely on geometric and physicochemical complementarity.

To balance the trade-off between docking and co-folding, DockFormer adopts a unique approach: while it co-folds the receptor, it requires the backbone distogram — representing C*α*-C*α* distances from a predicted or experimentally determined apo structure — as input. Since the backbone distogram serves only as an optional structural constraint, the model retains the ability to co-fold receptor-ligand complexes while avoiding the complexities of full protein folding. This design choice simplifies inference and reduces computational costs.

## 2 Methods

### 2.1 DockFormer Architecture and Key Modifications

The DockFormer architecture builds upon AlphaFold2 [13] architecture, with several key modifications to support ligand modeling and enhance efficiency. The most notable change is the replacement of MSA and template inputs with an input receptor structure. Rather than inferring contacts from evolutionary information in the MSA, the model directly receives approximated contact data. This modification simplifies the Evoformer, making it more similar to the Pairformer architecture introduced in AlphaFold3 [11]. The number of layers in the Evoformer portion of the model is reduced to 8 (instead of 48 in AlphaFold2). Another major adaptation is the incorporation of tokens representing ligand atoms alongside amino acid tokens. Lastly, DockFormer incorporates an affinity module, enabling multi-modal training for both docking and affinity prediction. To the best of our knowledge, this is the first adaptation of the AlphaFold2 architecture for both docking and affinity prediction.

The first step is the input embedding. The input for DockFormer is an AlphaFold2 prediction, or an experimentally determined apo conformation, of the receptor structure and a reference structure of the ligand generated by RDKit [33]. The output of the embedding stage is a single embedding and a pair embedding. The single embedding is a 1D vector containing an embedding vector for each token - either an amino acid or a ligand atom. The pair embedding is a 2D vector that contains an embedding for each pair of tokens. The features that are used for embedding protein tokens are only the amino acid types. For ligand tokens, the features used are atom types, charge, chirality, and bond types between atoms. Additionally, the input structures are converted into intra-protein and intra-ligand distograms, that is the distances between each pair of tokens. For proteins, C*α*-C*α* distances are discretized into 15 bins evenly distributed between 3.25Å and 20.75Å, similar to the recycling process in AlphaFold2. For ligands, the interatomic distances are assigned to one of the 10 bins, evenly distributed between 0.75Å and 9.75Å. The encoding of each matched bin is added to the embedding in the pair embedding.

After generating single and pair embedding, those are passed through a transformer-based model, with 8 blocks, where each block is similar to a Pairformer block. Recycling is also applied similarly to AlphaFold2 with 3 recycles. The final single- and pair- representations that are outputted by the last block are passed to the structure module as implemented in AlphaFold2. For ligand tokens, the frame rotations are ignored, and only transformations of frames are used as the atom positions. An additional optional step is to replace the predicted ligand positions with the closest conformation from a set of 1,000 conformations by RDkit. This step prevents impossible bond distances or conformations.

### 2.2 Affinity Prediction Module

In addition to the structure module, there is also an affinity module, which predicts the affinity as a classification task. The classification task is defined as 30 bins for affinities between 0 and 15, where the number represents the minus log of the nanomolar measured affinity. The final predicted affinity is a weighted average of the bins, weighted by the probability predicted for each bin. To predict the probability of each affinity bin, we test two approaches - the first is interface-based and the second is based on a classification token.

In the first approach, we apply a linear layer to every vector in the pair representation that represents a pair of a ligand atom and a receptor amino acid. Each such vector is weighted by a layer designated to weigh vectors for affinity relevance, and also by considering the logits predicted by the inter-contact auxiliary head. The weighted sum is used to predict the probability for each affinity bin.

In the second approach, we add a classification token to the model input, similar to what is common in Natural Language Processing models [34]. The vector corresponding to the affinity token in the single representation is passed through a linear layer to predict the affinity using regression. Although the interface-based approach is more complicated, it leverages data learned from structures, in the form of receptor-ligand contacts and encourages biological restraints on affinity prediction.

### 2.3 Training and model variants

We have implemented several new auxiliary heads and loss modifications. The inter-contact auxiliary head is trained to predict, given a pairwise representation, the binary classification task of whether the amino acid C*α* atom and ligand atom have a distance of at most 5Å. The binding site prediction head predicts for a single representation of a receptor token whether this token is within at most 5Å of the ligand. We have also added a variation of FAPE loss that considers all frames of the receptor, but only the ligand atoms, and by weighing this loss, we have forced the model to emphasize the contact between receptor and ligand.

We trained three versions of the model using different training datasets: DockFormer-PDBBind, DockFormer-Screen, and DockFormer-PLINDER.

**DockFormer-PDBBind** was trained on 18,101 samples from the PDBBind 2020 dataset. To enable fair evaluation against CASF-2016, we filtered out 479 samples from the original PDBBind dataset that were highly similar to CASF-2016, following the approach of previous methods [20]. Specifically, we removed any sample where the ligand’s ECFP4 fingerprint had a Tanimoto coefficient above 0.90 compared to any ligand in CASF-2016 or where a protein chain had over 90% sequence identity with a chain in CASF-2016.

**DockFormer-Screen** was designed to enhance the model’s ability to generalize for ligand screening, where a single protein interacts with many ligands. Since PDBBind contains diverse protein-ligand complexes, direct training on it does not optimize for this scenario. To address this, we curated a dataset of 12,374 samples across only 119 proteins (averaging 104 ligands per protein) and fine-tuned DockFormer-PDBBind on it. Affinity data was sourced from BindingDB [35], and receptor-ligand structures were predicted using Boltz-1 [36].

**DockFormer-PLINDER** was trained on the PLINDER [37] training set to assess generalization on its strictly separated test set. To maximize the utility of the dataset, we divided it using provided structural clusters and sampled data during training inversely proportional to cluster size. We excluded clusters representing different binding pockets of the same ligand (i.e., same protein and ligand but different structural clusters), as these could introduce ambiguity. Additionally, we capped the number of times each ligand could appear in the training set at five to prevent bias toward specific ligands. After filtering, we retained 6,979 clusters with a total of 138,402 samples. Finally, to enhance affinity prediction, we fine-tuned the model on all 36,594 affinity-labeled samples in the training set.

All models were trained with a batch size of 16, and after 25,000 optimization steps, we introduced an additional loss term on interatomic distances within the ligand to improve structural learning.

## 3 Results

### 3.1 CASF-2016 Affinity Benchmark

The CASF-2016 benchmark [38] is widely used to evaluate affinity prediction models, particularly those trained on PDBBind. It consists of 57 proteins, each with five ligands for which experimentally measured affinities are available. This benchmark allows for assessing a model’s ability to predict binding affinities and, separately, its ability to rank different ligands for the same protein, making it suitable for evaluating both general affinity prediction and ligand screening performance.

To assess model performance, we use different correlation metrics depending on the nature of the prediction task. Pearson correlation measures the linear relationship between predicted and experimental affinity values and is typically used when evaluating global affinity prediction, where receptor-ligand pairs are unrelated. However, when comparing screening performance, where multiple ligands must be ranked relative to a single protein, Kendall’s Tau is the preferred metric[39]. Kendall’s Tau assesses the consistency of ligand ranking, regardless of absolute affinity values, making it a more appropriate measure of a model’s ability to differentiate strong from weak binders.

Using CASF-2016, DockFormer-PDBBind achieved a Pearson correlation of 0.803 when using the interface-based affinity module, and 0.817 when using the classification-token-based affinity module (Fig. 2A). These results are comparable to state-of-the-art methods (Table 1).

**Table 1:** Comparison of affinity prediction performance across two benchmarks (CASF-2016 and PLINDER) using DockFormer and competing methods. Results are reported for two DockFormer affinity modules: interface-based (I) and token-based (T). Performance is evaluated using Root Mean Square Error (RMSE) and Pearson correlation. Lower RMSE and higher Pearson correlation indicate better performance.

| Method | CASF-2016 |  | PLINDER |  |
| --- | --- | --- | --- | --- |
|  | RMSE ↓ | Pearson correlation ↑ | RMSE ↓ | Pearson correlation ↑ |
| ConPlex[26] | - | - | - | 0.290 |
| Pafnucy[21] | 1.42 | 0.78 | 1.322 | 0.569 |
| PLANET[20] | 1.247 | 0.824 | 1.742 | 0.347 |
| DockFormer (I) | 1.299 | 0.803 | 1.199 | 0.656 |
| DockFormer (T) | 1.262 | 0.817 | 0.960 | 0.753 |

**Figure 1:**
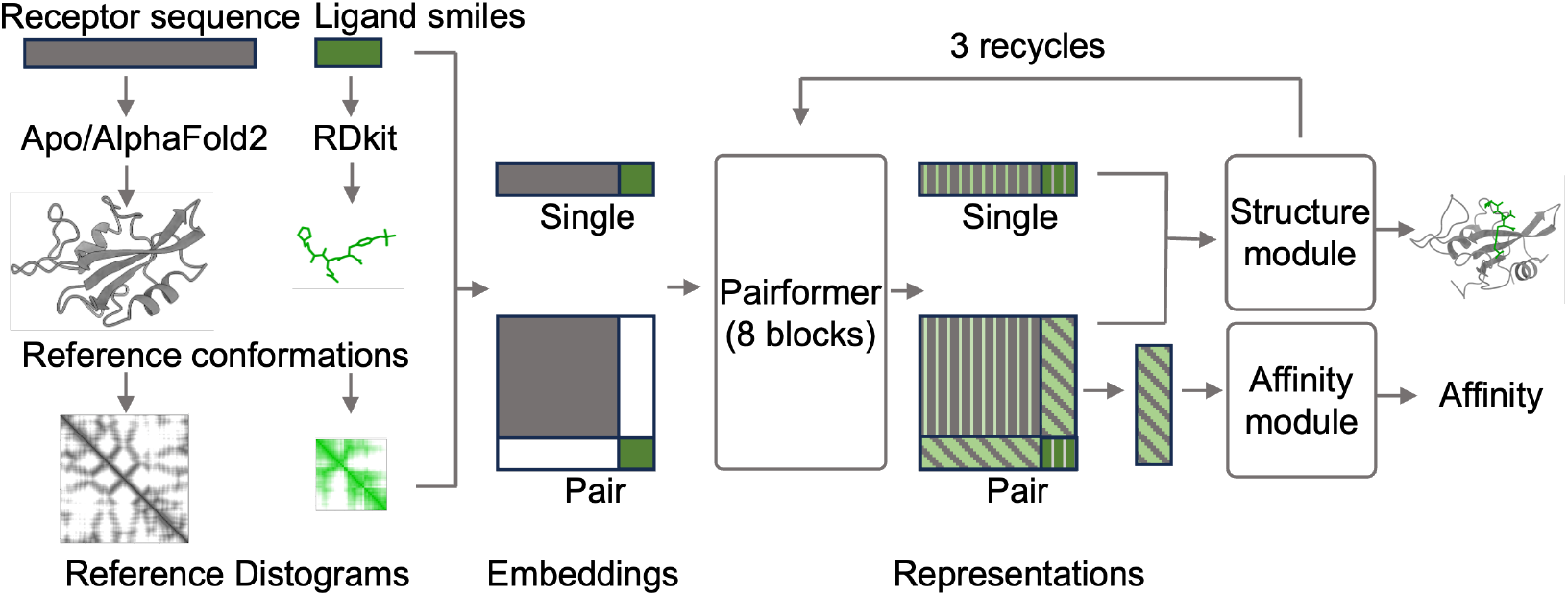
The model architecture. The inputs to DockFormer are receptor and ligand structure distograms. The structures are not required to be in the holo conformation, they can be an apo-conformation experimental structure or a predicted structure. For the ligand, a random reference conformation is generated. The distogram of the structures and the types of amino acids and atoms are then used to create a one-dimensional embedding for each input token referenced as single embedding and a two-dimensional input embedding for each pair of tokens referenced as pair embedding. These embeddings are the input for the deep learning-based model. The model consists of 8 Pairformer blocks that each output a single and pair representation. Both representations are used by the structure module to generate a structure. Only the parts in the pair representation that represent a pair of a ligand atom and a receptor amino acid are considered in the affinity module.

**Figure 2:**
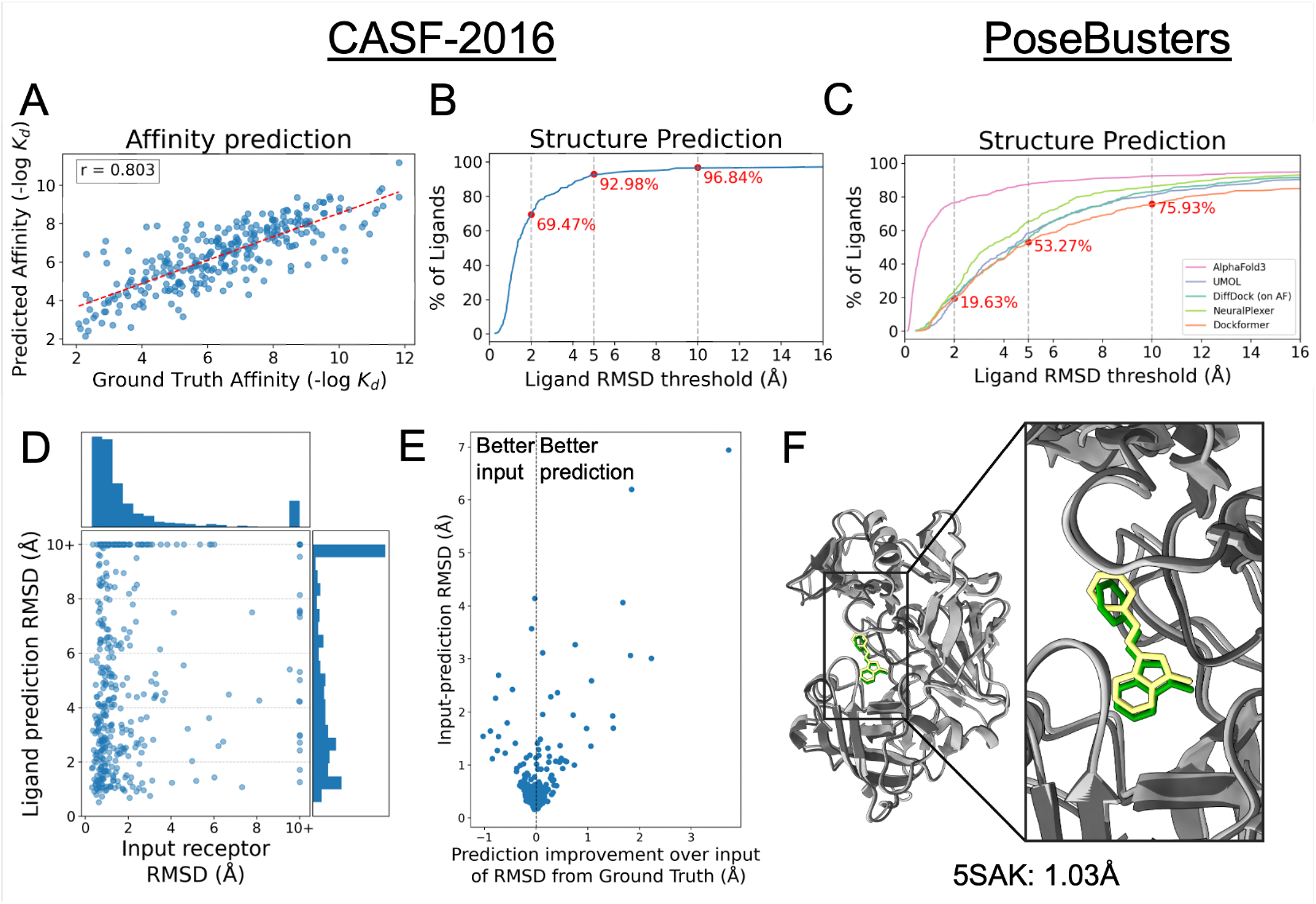
Results on CASF-2016 and PoseBusters benchmarks. **A** Predicted affinity compared to experimentally measured affinity in the CASF-2016 dataset (n=285). Axes are the minus log of the nanomolar Ki or Kd affinity. Pearson correlation of 0.803. **B** Structure prediction results for samples in the CASF-2016 dataset under continuous threshold. **C** Structure prediction results for samples in the PoseBusters dataset (n=428) under continuous threshold. **D** RMSD between predicted ligand positions when pockets aligned, compared to RMSD between ground truth receptor structure and input receptor structure. **E** The y-axis is the RMSD between the input receptor structure and the predicted receptor structure. The x-axis is the result of subtracting the RMSD between the input structure and the ground truth structure from the RMSD of the predicted structure and the ground truth structure. Positive x values represent an improvement in the accuracy of DockFormer prediction compared to the input structure. **F** Example of accurate prediction by DockFormer from the PoseBusters benchmark. Predicted receptor (light gray) and ligand (green) structure vs. experimental receptor (dark gray) and ligand (yellow) structure.

CASF-2016 can also be used as a ranking power benchmark, where Kendall’s Tau is computed separately for each of the 57 proteins by comparing the predicted rankings of their five ligands. Using this metric, DockFormer-PDBBind achieved an average Kendall’s Tau of 0.60, while DockFormer-Screen improved this to 0.62. This suggests that fine-tuning on screening-specific data enhances the model’s ability to rank ligands within the same protein, as opposed to a model trained solely on general receptor-ligand affinity predictions. These results highlight DockFormer’s strong performance in both global affinity prediction and ligand screening, reinforcing its effectiveness in structure-based drug discovery applications.

### 3.2 Structure Prediction on CASF-2016 and PoseBusters

For structural evaluation, we assessed ligand Root Mean Square Deviation (RMSD) relative to the experimental structure, aligning based on the C*α* atoms of the protein pocket amino acids - amino acids with at least one atom within 8Å of any of the ligand atoms. A ligand RMSD below 2Å is considered a successful prediction, while an RMSD under 10Å generally indicates correct binding pocket identification. DockFormer accurately predicted ligand structures with RMSD < 2Å for 69% of complexes in the CASF-2016 dataset and correctly identified the binding pocket (RMSD < 10Å) for 93% of complexes (Fig. 2B).

Despite filtering the training data to remove complexes similar to those in CASF-2016, it appears that the dataset still contains structurally related complexes. To further evaluate the generalization ability of DockFormer for structural prediction, we used PoseBusters, a widely adopted benchmark for protein-small molecule interactions [40]. This dataset consists of 428 interactions, none of which are included in PDBBind. On this benchmark, DockFormer-PDBBind successfully predicted 20% of complexes with ligand RMSD <2Å, a performance comparable to competing methods [10, 16, 41], except for AlphaFold3 and its variants [11, 36, 42] (Fig. 2C,F).

A particularly interesting comparison is between DockFormer and Umol [16], where DockFormer achieved 20% success, slightly outperforming Umol’s 18% success rate. Since both models are based on the AlphaFold2 architecture, the main distinctions are DockFormer’s reduced number of layers and its removal of MSA from the pipeline. This suggests that using input structures instead of co-folding from scratch can maintain performance while significantly reducing model complexity.

### 3.3 Impact of Input Receptor Structure Accuracy

To analyze the relationship between input receptor structure accuracy and docking performance, we examined the C*α* RMSD between the input and ground truth structures in the PoseBusters test set. In 73% of cases, the input receptor structure had C*α* RMSD <2Å, and only 8% of samples exceeded 10Å RMSD (Fig. 2D).

We also investigated protein flexibility during docking by comparing the input and output protein structures. Excluding 38 cases where the input structure was either highly inaccurate (>10Å RMSD) or too large (>1,300 residues), we found that in 97% of cases, the receptor structure remained largely unchanged (C*α* RMSD <1Å, Fig. 2E). However, in cases where significant changes occurred, the predicted structure was often closer to the ground truth than the input, suggesting that DockFormer refines receptor structures to better fit the ligand.

To test whether DockFormer depends on an accurate receptor conformation, we evaluated its performance using holo-input structures instead of AlphaFold2-predicted receptor structures. This resulted in a slight performance improvement from 20% to 21%, indicating that DockFormer is not strongly dependent on receptor structure accuracy.

### 3.4 Generalization on PLINDER

It has been observed that structure prediction accuracy correlates with the similarity between test samples and training data [40]. The PLINDER dataset [37] was recently introduced with stricter train-test separation criteria to address this concern. To benchmark generalization, we trained DockFormer-PLINDER exclusively on the PLINDER training set and tested it on the PLINDER test set.

On PLINDER, DockFormer’s structure prediction success rate (ligand RMSD <2Å) dropped from 20% to 15% compared to PDBBind training. This highlights the importance of avoiding data leakage while also demonstrating DockFormer’s ability to generalize.

For affinity prediction, we evaluated 112 test set samples with experimentally measured affinities. Using the interface-based affinity module, DockFormer achieved a Pearson correlation of 0.656, which increased to 0.753 when using the classification-token affinity module. This aligns with the minor drop in structure prediction accuracy, while still maintaining strong overall performance. Notably, competing methods achieved a maximum Pearson correlation of 0.569 (Table 1).

Additionally, some of these competing methods [20, 21] require the experimental bound structure to predict affinity, whereas DockFormer operates fully independently. Furthermore, all competing methods were trained on datasets without strict train-test separation, raising the possibility of data leakage.

### 3.5 Validation on CASP16 ligand targets

To further validate DockFormer, we evaluated its performance in the CASP16 Pharmaceutical Protein-Ligand Challenge, which includes two target proteins (Chymase and Autotaxin) with multiple ligands and experimentally measured affinities. We compared DockFormer-Screen to the top-performing groups in the challenge for both structure and affinity prediction, using the interface-based affinity module for affinity estimation.

DockFormer’s results were comparable to the top-scoring groups in both structure and affinity prediction (Fig. 3). Notably, while most CASP16 participants utilized templates and manual refinements, DockFormer generated fully automated predictions with no manual intervention. It is also important to emphasize that we compare against only the top-performing groups among at least 32 competitors, representing state-of-the-art methods in receptor-ligand interaction prediction.

**Figure 3:**
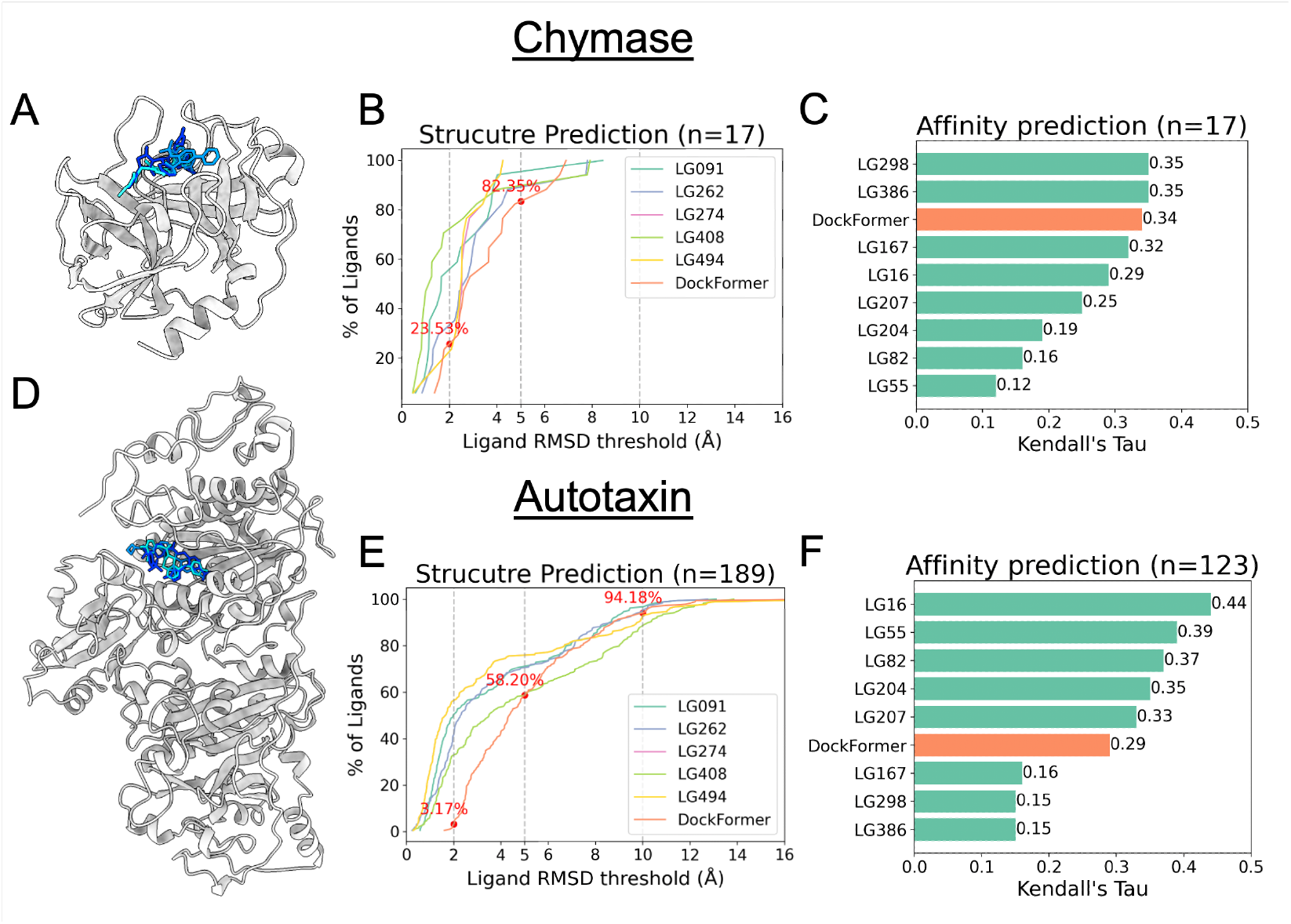
Results on CASP16 benchmark. **A** Experimental structure of Chymase (gray) with 3 of the ligands imposed onto it (shades of blue). **B** Structure prediction accuracy for 17 ligands of Chymase structure under a continuous RMSD threshold. Accuracy is shown for DockFormer and for top-performing groups in structure prediction of ligands in CASP16. **C** Kendall’s Tau of DockFormer-Screen interface-based affinity predictions and experimentally measured affinities for 17 ligands bound to Chymase. Compared to top-performing groups in affinity prediction in CASP16. **D** Experimental structure of Autotaxin (gray) with 3 of the ligands imposed onto it (shades of blue). **E** Structure prediction accuracy for 189 ligands of Autotaxin structure under a continuous RMSD threshold. Accuracy is shown for DockFormer and for top-performing groups in structure prediction of ligands in CASP16. **F** Kendall’s Tau of DockFormer-Screen interface-based affinity predictions and experimentally measured affinities for 123 ligands bound to Autotaxin, 93 of them also had structure. Compared to top-performing groups in affinity prediction in CASP16.

When comparing DockFormer-PDBBind to DockFormer-Screen, we observe no substantial difference in structure prediction, but a clear improvement in affinity prediction for DockFormer-Screen. For Chymase, DockFormer-PDBBind achieves a Kendall’s Tau of 0.24 (compared to 0.34 with DockFormer-Screen), while for Autotaxin, the correlation is 0.18 (compared to 0.29 with DockFormer-Screen). In contrast, DockFormer-PLINDER performs significantly worse in both structure and affinity prediction, achieving only 0.04 for Chymase and 0.05 for Autotaxin in affinity correlation.

Additionally, we compared the classification-token-based affinity module to the interface-based module. For Autotaxin, the classification-token module yielded slightly better performance (0.30 vs 0.29), whereas for Chymase, performance dropped significantly to (0.06 vs.0.34), highlighting the model’s sensitivity to specific proteins, suggesting that certain affinity modules may be more suited to particular receptor-ligand interactions.

### 3.6 Further analysis

A key strength of DockFormer is its lightweight architecture, enhancing prediction efficiency. On the PoseBusters dataset, the median prediction time was 2.67 seconds, with an average of 6.11 seconds. Although these run times do not account for the generation of the input receptor structure, they remain highly relevant for screening processes where the same receptor is tested against multiple ligands, making the receptor structure generation a negligible factor.

One could expect that the accurate prediction of affinity will be dependent on the accurate prediction of structure, as affinity is highly dependent on the protein-ligand contacts. However, it is clear that in many cases DockFormer was not able to predict an accurate structure, but was able to predict an accurate affinity. This was also seen during training, where the affinity prediction loss converged much faster compared to structure prediction losses. This is not completely non-intuitive, as there are many known models for affinity prediction that are not based on structure. This shows that the Pairformer architecture can learn to predict the interaction properties, even without predicting the structure itself.

## 4 Discussion

We have presented DockFormer, a method for predicting structure and affinity of receptor-ligand interactions that integrates two key concepts into the AlphaFold2 [13] architecture. First, the method uses an approximated receptor structure, either apo conformation or prediction, as an input, enabling the model to specialize in the docking task over the protein folding task, while keeping the receptor structure flexible to capture conformational changes. This also enables reducing the model architecture to provide efficiency in training and inference. Second, by employing multi-task learning for simultaneous affinity and structure prediction, the network gains additional information that can further enhance its accuracy.

Our results demonstrate that DockFormer achieves competitive structure prediction performance while excelling in affinity prediction and ligand ranking. While DockFormer successfully predicted ligand poses with reasonable RMSD values, its structural accuracy remains lower than that of diffusion-based models such as AlphaFold3 [11] and Boltz-1 [36]. One potential reason for this difference is that diffusion architectures explicitly model fine-grained ligand-receptor interactions and optimize side-chain flexibility, whereas DockFormer treats side-chains implicitly, which may limit structural accuracy. Despite this, DockFormer demonstrated strong generalization on the PoseBusters and PLINDER benchmarks while maintaining high efficiency, making it well-suited for high-throughput virtual screening where speed is crucial.

Beyond structure prediction, DockFormer demonstrated state-of-the-art affinity prediction performance, achieving high Pearson correlations across multiple benchmarks, including CASF-2016 and PLINDER. Its ability to rank ligands within the same protein, as seen in CASF-2016 scoring benchmarks and CASP16, further underscores its effectiveness in screening scenarios. Interestingly, even in cases where structural accuracy was limited, DockFormer often produced accurate affinity predictions, suggesting that the Pairformer architecture effectively captures binding interactions without relying on perfect structure prediction. These findings suggest that DockFormer is a powerful and scalable approach to structure-based drug discovery, with potential for further refinement through architectural enhancements, such as diffusion-based modeling, to improve structural accuracy while maintaining its efficiency and affinity prediction accuracy.

The key concepts introduced here — leveraging input structures and integrating affinity modules — can be applied to emerging high-performance architectures. These approaches can be implemented either by training a new, streamlined model with fewer layers to improve efficiency or by fine-tuning existing models to incorporate these capabilities while benefiting from their already well-optimized weights. In future work, we plan to explore both strategies and evaluate their impact on accuracy.

The foundational concepts behind DockFormer have the potential to be extended beyond receptorligand interactions. For instance, similar models could be developed for protein-protein interactions, including specialized models for challenging cases like antibody-antigen binding or protein-peptide interactions. In those cases, the additional modality of the model can be specificity. By leveraging large databases such as STRING[43], which provide interaction labels, we can train models to predict whether proteins interact. The reduction DockFormer offers in model size may make specialized task-specific models more feasible. We provide below an initial implementation of protein-protein interaction prediction named DockFormerPP, which we intend to analyze and expand in future work.

## 5 Code Availability

DockFormer was implemented by building upon the OpenFold[44] implementation of AlphaFold2[13]. The code for running and training DockFormer, with the weights of three presented variations is available as code and as a server at: https://huggingface.co/spaces/benshor/DockFormer.

Additionally, the variation DockFormer-PINDER for protein-protein interaction prediction is available at: https://huggingface.co/spaces/benshor/pinder_dockformer/tree/main.

